# Chromosome-level genome construction of a Japanese stickleback species using ultra-dense linkage analysis using single-cell sequencing of sperms

**DOI:** 10.1101/2020.05.12.092221

**Authors:** Kazutoshi Yoshitake, Asano Ishikawa, Ryo Yonezawa, Shigeharu Kinoshita, Jun Kitano, Shuichi Asakawa

**Author notes:** Corresponding author: Shuichi Asakawa.

## Abstract

The presence of high quality genomes at the chromosome level is very useful in the search for the causal genes of mutants and in genetic breeding. The advent of next-generation sequencers has made it easier to decode genomes, but it is still difficult to construct the genomes of higher organisms. In order to construct the genome of a higher organism, the genome sequence of the organism is extended to the length of the chromosome by linkage analysis after assembly and scaffolding. However, in the past linkage analysis, it was difficult to make a high-density linkage map, and it was not possible to analyze organisms without an established breeding system. As an innovative alternative to conventional linkage analysis, we devised a method for genotyping sperm using 10x single-cell genome (CNV) sequencing libraries to generate a linkage map without interbreeding individuals. The genome was constructed using sperm from *Gasterosteus nipponicus*, and single-cell genotyping yielded 1,864,430 very dense hetero-SNPs. The average coverage per sperm cell is 0.13x. The number of sperm used is 1,738, which is an order of magnitude higher than the number of sperm used for conventional linkage analysis. We have improved the linkage analysis tool SELDLA (Scaffold Extender with Low Depth Linkage Analysis) so that we can analyze the data in accordance with the characteristics of single-cell genotyping data. Finally, we were able to determine the location and orientation on the chromosome for 85.6% of the contigs in the 456 Mbase genome of *Gasterosteus nipponicus* sequenced in nanopores. A total of 95.6% of the contigs in which a cross-reaction was detected within the contigs.

## Introduction

The development of next-generation sequencers (NGS) and their peripheral technologies has provided an environment for easy genome decoding. In particular, it is possible to decode a bacterial genome with a genome size of several million bases by using nanopores and PacBio (Liao et al., 2015; Wick et al., 2017). However, the genome size of higher eukaryotes is often more than 100 times larger than that of bacteria, and there are large repetitive sequences such as centromeres, telomeres, and clusters of rRNAs, making it difficult to decipher the genomes of higher eukaryotes. In the genome decoding process, after assembling the short sequencing data and scaffolding the assembled contigs, it is extended to the length of the chromosomes through linkage analysis or Hi-C. However, the step from scaffolds to chromosomes remains challenging. Of the 9,363 eukaryotic species in the NCBI genome database, only 16% have been extended to chromosomes, and many organisms have not constructed chromosome-level genomes yet. Each of the analysis methods used in this chromosome elongation step has problems. In linkage analysis, it is necessary to prepare the DNA of the parents and children in order to detect the recombination that occurs in the germ line cells of the parents. However, it is often difficult to prepare the DNA of the parents and children in many species whose genomes are still undetermined because the breeding system has not been established. Meanwhile, the recently developed Hi-C method, which utilizes the higher-order structure of the genome in the cell nucleus, can be analyzed by only a single individual and can detect genomic interactions with a resolution of 1 kb (Rao et al., 2014), but there is uncertainty in terms of accuracy because there is no information on the distance between the interacting regions.

We have developed methods to construct genomes at the chromosome level using linkage analysis, such as linkage analysis using female monozygotic individuals (doubled haploid) (Fig. 1B) (Zhang et al., 2018) and linkage analysis using hybrids (Yoshitake et al., 2018). A common point of focus for these methods is that genotyping a diploid genome usually requires 30x or more coeverages, whereas a haploid genome only needs to be read 1x. However, it was possible to improve the resolution of the markers significantly, and make a very accurate linkage map of any species that has an established breeding system in exchange for more restrictions than the traditional linkage analysis. In this study, we further developed the doubled-haploid and cross-species system and incorporated the recently developed single-cell technology for genotyping of single sperm cells, where each cell is assigned a different barcode sequence in a 10x Genomics library conditioning system, which is read at once by an Illumina sequencer, and the data is separated for each cell by barcode during data analysis (Fig. 1C). Therefore, it is possible to easily sequence even 10,000 cells by increasing the amount of reads. If sperm can be genotyped in a single cell, it is possible to directly detect recombinations that occur in the germ line cells of the parent, without the need to prepare their offspring. Furthermore, since sperm is a haploid, it is possible to efficiently genotype and a densely linkage map from the 1x and smaller data volume cultivated in our doubled hapl oids and hybrids systems. Genotyping from single sperm cells has been reported in humans, Daphnia, and others (Xu et al., 2015; Zong et al., 2012), but the number of cells analyzed is as low as 104 at best. Using the 10x single cell system, it is realistically possible to analyze 10,000 or more sperms. In human, if 10,000 individuals can be used, the detectable recombination reaction will be once about every 10 kb, because the genetic distance is roughly 1 cM = 1,000 Kb, which will result in one recombination in every 100 individuals per 1,000 Kb (Broman et al., 1998). If a recombination reaction occurs in a scaffold, the position and orientation of the scaffold in the final chromosome can be completely determined. In the past, the size of the scaffold used for linkage analysis was usually required to be a megabase class because of the small number of markers and the small number of individuals to be used. In addition, conventional linkage analysis requires only 100 individuals and at most 10,000 markers, so analysis tools can not handle 10,000 individuals and 1 million markers. We have developed a software called SELDLA as a tool for ultra-high-density linkage analysis in doubled-haploid and crossbreeding species, and by improving SELDLA to apply the data of single-cell sperm systems. SELDLA constructed a genome that makes efficient use of single-cell data. In this study, single-cell whole genome amplification is performed on sperms of *Gasterosteus nipponicus* using 10x single-cell genome (CNV) sequencing libraries. The genome size of *Gasterosteus nipponicus* is 465 Mb, which is relatively small among vertebrates and has been the focus of attention in the study of freshwater penetration of fishes (Ishikawa et al., 2019; Yoshida et al., 2016) and is a closely related species of the extensively studied three-spined stickleback (Gasterosteus aculeatus) (Jones et al., 2012; Kingsley and Peichel, 2006).

**Figure 1.**
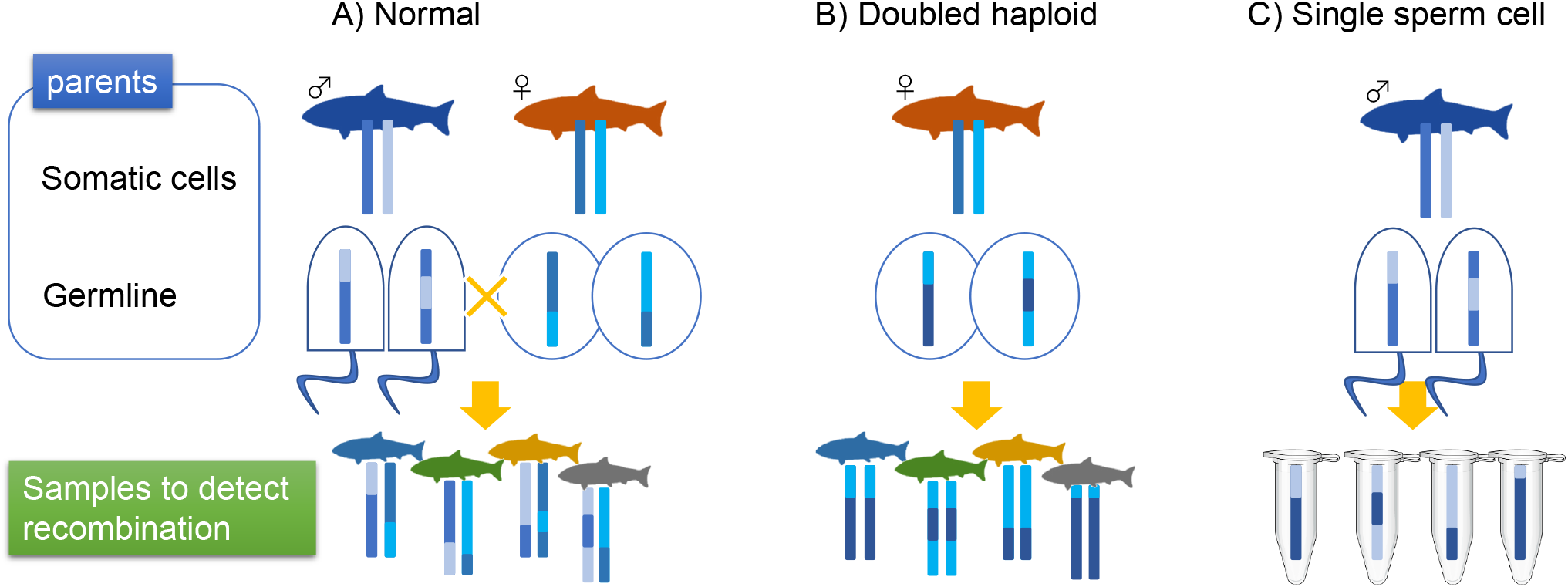
Comparison of linkage analysis methods A) Normal linkage analysis. The father and mother each have a diploid genome, and the children inherit two of the parent’s four total haplotypes. The genome of a hetero-diploid children will be used to detect recombination. b) Linkage analysis using doubled haploid. Recombination is detected in children that are produced by artificial parthenogenesis from the mother’s oocyte. The children have a homozygous diploid genome. c) Linkage analysis using single sperm cells. Recombination is detected in the haploid single sperm after whole-genome amplification.

## Results and Discussions

### - Single-cell library construction

Sperms were collected from an individual of *Gasterosteus nipponicus* captured at Akkeshi Bay and a single-cell library was constructed by 10x chromium using approximately 300,000 sperms. The constructed libraries were sequenced by HiSeq. The number of sequenced reads was 1,276,740,950 x 2 paired-end reads with a total base of 383 Gb.

### - Nanopore sequencing

DNA was extracted from the liver and fin + pectoral fin muscles of the individual from which sperm was collected, and the length of the DNA fragments was confirmed by electrophoresis. The length of the extracted DNA from fin + pectoral fin muscles was longer than that from liver. The length of the DNA fragment of the fin + pectoral fin muscle was checked by a TapeStation (Figure 2A), and because the 7-60 Kb long fragment was only 57.95% (Figure 2B), short reads were removed with the Short Read Eliminator. As a result, a long fragment of 7-60 Kb was enriched to 76.54% (Figure 2C). The number of reads obtained by nanopore sequencing was 2,163,108, with a total base count of 8.1 Gb (17.8x) and a read N50 of 5.87 Kb. The sequenced reads were assembled using Flye. The number of contigs obtained was 3,895, the N50 was 0.563 Mb, and the BUSCO score was 81.2% (Table 1). The resulting contigs were polished using nanopore reads and single-cell HiSeq reads, and the post-polishing genome showed an elevated BUSCO score of 96.7% (Table 1), confirming that the assembly error could be corrected.

**Figure 2.**
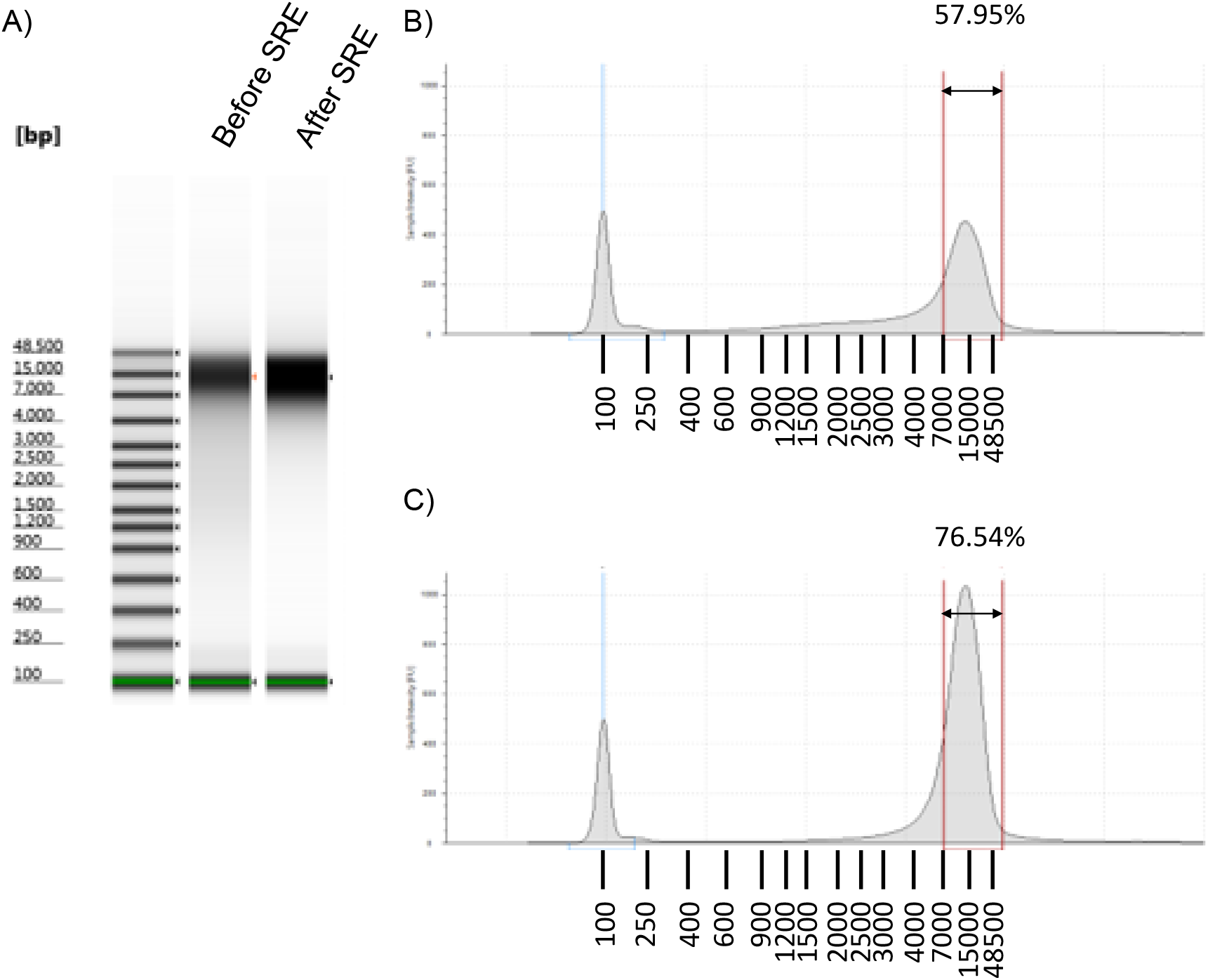
Extracted DNA length A) Results of electrophoresis using a TapeStation. B) Densitometry of lectrophoresis using a TapeStation before SRE treatment. 100 bp peak is a marker. The percentage of DNA fragments in the range of 7,000 - 60,000 is shown in the figure. C) Densitometry using a TapeStation after SRE treatment.

**Table 1.** Statistical information of the assembled genome and the polished genome

|  | **N** | **N50 [Mb]** | **Total [Mb]** | **BUSCO [%]** |
| --- | --- | --- | --- | --- |
| Assembled | 3,895 | 0.563 | 455 | 81.2 |
| Polished | 3,668 | 0.565 | 456 | 96.7 |

### - Genotyping single sperm cells

The analysis of the sequenced data read with 10x single-cell genome (CNV) sequencing libraries was analyzed using Cell Ranger software to detect copy number variations of each cell with the polished genome as a reference genome. CNV analysis resulted in the detection of 2,286 cells (Table 2). However, because barcodes with a high number of reads contain multiple cells, we removed the cells that were judged to be more than doubles by the score of ploidy_confidence. As a result, 1,738 cells remained. A total of 1,864,430 heterozygous SNPs were detected in these cells. This means that there is one SNP in every 245 bases because the genome size is 456 Mb. The average coverage per cell was 0.13x.

**Table 2.** The result of 10x single-cell genome (CNV) sequencing

|  |  |
| --- | --- |
| The number of sequenced reads | 1,276,740,950 x 2 |
| The sequenced bases | 383,022,285,000 bases |
| The number of sequenced cells | 2,286 |
| The number of haplotype cells | 1,738 |
| The number of total heterozygous SNPs | 1,864,430 |
| The average coverage per cell | 0.13x |

### - Genome construction from single sperm cells

The extracted SNPs were input to SELDLA for the linkage analysis. This resulted in 21 scaffolds of 10 Mb or more, consistent with the number of chromosomes in the closely related species *Gasterosteus aculeatus* (Figure 3). The SELDLA-extended scaffold N50 became 18.39 Mb (Table 3). The percentage of contigs that could be placed on 21 chromosomes was 87.2%, and the percentage of contigs that could also determine orientation was 85.6%. The percentage of bases from the contigs for which recombinations were observed was 95.6%. The contigs with recombination have the potential to determine their position and orientation in the genome. Future improvements of the SELDLA algorithm could theoretically place 95.6% of the contigs on the chromosome.

**Figure 3.**
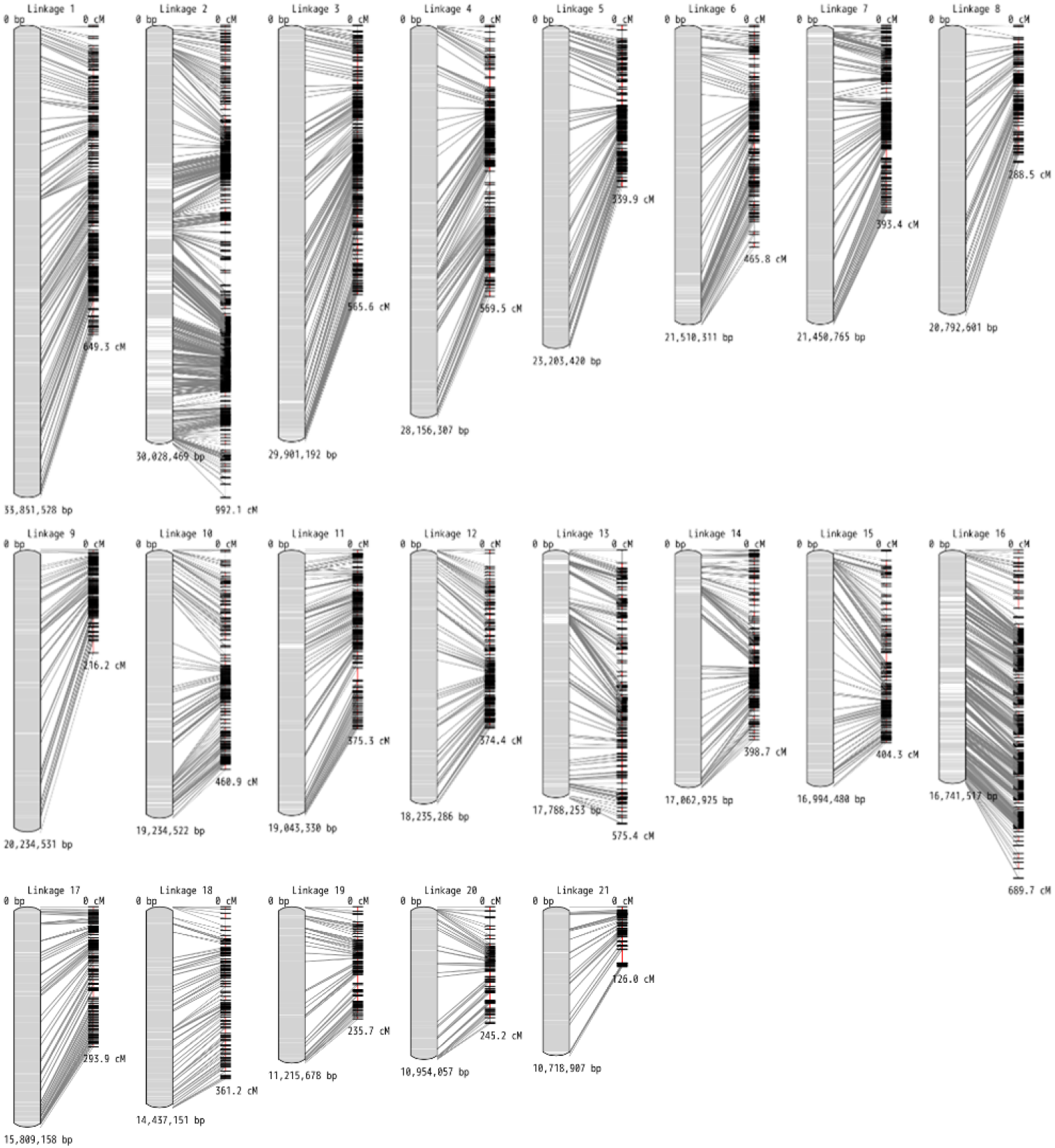
The physical map and the linkage map of the constructed genome by single sperm cell linkage analysis The correspondence between physical (left) and genetic (right) distances is shown for scaffolds elongated to more than 10 Mb. The lines indicating correspondence between physical and linkage maps are based on positions of both ends of the contigs.

**Table 3.** Statistical information of the SELDLA-extended genome

|  | **N** | **N50 [Mb]** | **Total**<br>**[Mb]** | **Placed bases in**<br>**chromosomes [%]** | **Oriented bases in**<br>**chromosomes [%]** | **BUSCO**<br>**[%]** |
| --- | --- | --- | --- | --- | --- | --- |
| Polished | 3,668 | 0.565 | 456 | - | - | 96.7 |
| SELDLA | 1,480 | 18.39 | 456 | 87.2 | 85.6 | 97.1 |

### - Comparison of the constructed genome with closely related species

Comparison with the genome of a closely related species, *Gasterosteus aculeatus*, was performed. The dot plots show that the published genome of *Gasterosteus aculeatus* and the genome of *Gasterosteus nipponicus*, which was generated in this study, roughly correspond to each other (Figure 4). In particular, chromosome 8 and chromosome 13 from *Gasterosteus aculeatus* and scaffold 7 and scaffold 6 from *Gasterosteus nipponicus* were shown to be highly homologous from end to end of the chromosome. On the other hand, for scaffold 16, which is homologous to Chromosome 19 of *Gasterosteus aculeatus*, there is local homology, but the homologous regions are not aligned on the diagonal. The sex determination type of *Gasterosteus nipponicus* is known as a XY type, and the sex chromosome of *Gasterosteus aculeatus* is known to be chr19. In this study, the linkage map was constructed from male sperm, there was n o recombination between the sex chromosomes in *Gasterosteus nipponicus*. It was the cause of the inability to accurately place the contigs because the linkage map of the sex chromosomes could not be made. The elongation of the sex chromosome is an issue that needs to be addressed in the future, given the use of sperm-based linkage maps. However, even in the present study, it was possible to recognize the sex chromosome-derived contigs as being derived from a single chromosome. In addition, the XY chromosomes differ by only one gene in medaka (Matsuda et al., 2002) and pufferfish (Kamiya et al., 2012), and crosses occur between the XY chromosomes in above species. Therefore, it is possible to fully determine the whole genome of organisms with poorly differentiated sex chromosomes using this single sperm cell method.

**Figure 4.**
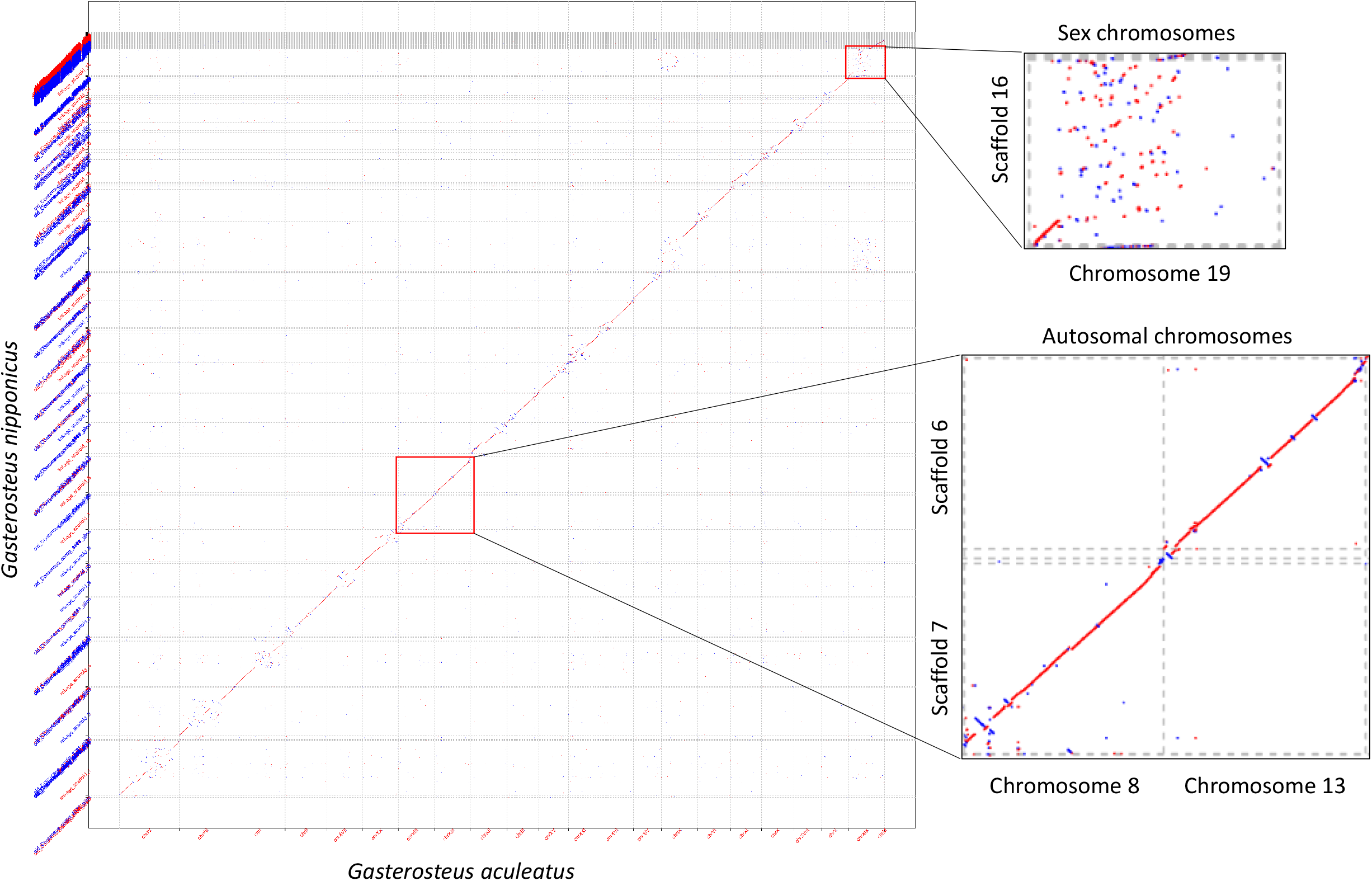
The whole genome dot plot analysis of *Gasterosteus nipponicus* and *Gasterosteus aculeatus* A dot plot comparing the SELDLA-elongated *Gasterosteus nipponicus* chromosome (Y axis) and the published *Gasterosteus aculeatus* reference sequence (X axis). Red lines in the dot plot indicate homology in the forward direction, and blue lines indicate homology in the reverse direction.

## Conclusion

This work established an innovative genome construction method for genotyping sperm using 10x single-cell genome (CNV) sequencing libraries and generating linkage maps without interbreeding. The detected SNP markers were very dense (1,864,430) and the number of sperm that could be analyzed was 1,738, which is an order higher than the number of individuals in conventional linkage analysis. Theoretically, more than 95% of the contigs can be placed on chromosomes, and the genome can be constructed with great accuracy. The linkage analysis method using single sperm cells is very versatile and is expected to become a standard method for genome analysis in the future.

## Methods

### - Sperm collection from *Gasterosteus nipponicus*, DNA extraction from liver, fin and pectoral muscles

The G.nipponicus fish was a gift from a local fisherman, who caught the fish in Akkeshi Bay, Hokkaido, Japan in May 2019. After collection, the fish was maintained in 50% seawater at 16°C under a long-day condition (16L8D) for 6 months. The fish was killed using an overdose of buffered MS222 (300 mg L-1). Male testes of *Gasterosteus nipponicus* were excised, the testes were disrupted in 200 *μ*l of cell suspension buffer (L15: FPS = 9: 1), sperm were released, and the sperm were well dispersed by pipetting. After removing the cell mass with a Cell strainer, the cell suspension was diluted to 1/100, and the cell count was done using an automated cell counter. Cell suspensions of sperm were centrifuged at 500 x g, 4°C for 5 min, then the supernatant was removed and resuspended in 400 *μ*l of cryopreservation solution (FPS: DMF = 9: 1), placed in Cryotube, lowered in temperature for 20 min in a 15 ml tube spiked with dry ice, and stored in liquid nitrogen. Other body parts were preserved in 100% Ethanol for DNA extraction. All animal experiments were approved by the institutional animal care and use committee of the National Institute of Genetics (31-16). DNA extraction from the liver or fin and pectoral fin muscles of the fish from which sperm was collected were performed using DNeasy Blood & Tissue Kit (QIAGEN).

### - Nanopore sequencing

To remove short fragments less than 7 Kb, Short Read Eliminator XS (SRE) (Circulomics) was used. The procedure was basically the same as the manufacturer’s protocol, but the temperature of elution was changed to 50°C using 50 *μ*l of Buffer EB. The amount of DNA after SRE elution was measured using Qubit 2.0 (Thermo Fisher Scientific) and Qubit dsDNA BR Assay Kit (Thermo Fisher Scientific) for a total of 5.5 *μ*g. To determine the length of DNA fragments after SRE, gDNA before SRE treatment was diluted 5-fold and the undiluted solution after SRE treatment was used according to the protocol using TapeStation 2200 (Agilent Technologie) and Genomic DNA ScreenTape (Agilent Technologie).

The amount of DNA at the start was adjusted to approximately 1.2 ug, and the library was prepared according to the protocol using the Ligation Sequencing Kit SQK-LSK109 (Nanopore). All DNA concentration measurements were performed using the Qubit 2.0 and Qubit dsDNA BR Assay Kit. Using Flow Cell R9.4.1 (Nanopore), the flow cell was washed twice and sequenced a total of three times; the number of active pores, initial bias voltage and sequencing times of the three nanopores were as follows. 1st run: 1,432, −180 mV, 13 h 57 m, 2nd run: 1,136, −190 mV, 19 h 48 m, 3rd run: 901, −210 mV, 72 h 8 m. For the flow cell wash, the EXP-WSH003 was used, and the reaction time of the WashMix was changed from 30 minutes to 2 hours and followed the protocol for the rest of the process. The base call was done by MinION Software 19.12.5.

### - Sperm single cell analysis

The frozen sperm were sent to GENEWIZ for 10x single-cell CNV library preparation and sequencing. Basically, the 10x Genomics Chromium single-cell CNV library preparation was performed according to the previous paper of single sperm cell analysis (Tomoiaga et al., 2020). The sequence was performed in 3 lanes with HiSeq 4000 with paired-end sequencing of 150 bp.

### - Creating contigs

The nanopore sequenced data were assembled using Flye (ver 2.7) (Kolmogorov et al., 2019) with the options --nano-raw, --genome-size 400M and --threads 12. For the assembled contigs, we mapped the nanopore sequence data with minimap (ver 0.2) (Li, 2016) and polished them with Racon (v0.5.0) (Vaser et al., 2017). 500,000,000 reads (311 folds of the genome) were extracted from single-cell Illumina data, 16 bp on the forward side was removed as it was a 10x cell barcode, mapped to the genome after polishing by racon with BWA MEM (ver 0.7.15) (Li, 2013), and polishing with Illumina data with Pilon (ver 1.23) (Walker et al., 2014).

### - SNP extraction

A single-cell CNV analysis was performed using Cell Ranger (ver 3.0.2) (10x Genomics) using the genome after pilon polishing as a reference sequence. Cell Ranger does not consider contigs less than 1 Mb in length, so pseudo chromosomes with short contigs connected by N x 10,000 were created and analyzed. We also used vartrix (v1.1.14) (10x Genomics) to call SNPs in single cells. Bcftools mpileup (ver 1.10.2) (Li, 2011) was used in advance to create a bulk vcf file of all SNPs from 500,000,000 reads from single cells and the bulk vcf file was inputted to vartrix. The matrix file of SNPs obtained from vartrix was converted to cell-separated vcf format by in-house script.

### - Genome elongation by linkage analysis

Called cell-separated vcf files were analyzed by SELDLA (ver 2.0.9) (Yoshitake et al., 2018) for the extension of contigs to chromosomes with the following options, --mode=haploid -p 0.03 -b 0.03 --cs 2 --nl 0.9 --NonZeroSampleRate=0.05 --NonZeroPhaseRate=0.1 -r 4000 --RateOfNotNASNP=0.001 --RateOfNotNALD=0.01.

### - Comparative genomics

We used minimap2 (Li, 2018) to compare homologous regions between *Gasterosteus aculeatus* (Glazer et al., 2015) and *Gasterosteus nipponicus*. The script is available as the script name of post-assemble~dotplot-by-minimap2 in a portable pipeline software (https://github.com/c2997108/OpenPortablePipeline).

## Author contributions

K.Y., J.K. and S.A designed the project. R.Y. and A.I. performed all wet-lab experiments. K.Y. performed all data analysis. K.Y., A.I., S.K. and S.A. wrote the manuscript with input from all authors. All authors read and approved the final manuscript.

## Data availability

All sequencing data is available in DDBJ under accession number “PRJDB9841”.

## Code availability

SELDLA software is available at https://c29979108.wixsite.com/seldla.

## Acknowledgements

This work was supported by JSPS KAKENHI Grant Number 20K06607. Computations were partially performed on the NIG supercomputer at ROIS National Institute of Genetics.

